# Stress alters social behavior and sensitivity to pharmacological activation of kappa opioid receptors in an age-specific manner in Sprague Dawley rats

**DOI:** 10.1101/399758

**Authors:** Elena I. Varlinskaya, Linda Patia Spear, Marvin R. Diaz

## Abstract

The dynorphin/kappa (DYN/KOR) system has been identified as a primary target of stress due to behavioral effects, such as dysphoria, aversion, and anxiety-like alterations that result from activation of this system. Numerous adaptations in the DYN/KOR system have also been identified in response to stress. However, whereas most studies examining the function of the DYN/KOR system have been conducted in adults, there is growing evidence suggesting that this system is ontogenetically regulated. Likewise, the outcome of exposure to stress also differs across ontogeny. Based on these developmental similarities, the objective of this study was to systematically test effects of a selective KOR agonist, U62066, on various aspects of social behavior across ontogeny in non-stressed male and female rats as well as in males and females with a prior history of repeated exposure to restraint (90 min/day, 5 exposures). We found that the social consequences of repeated restraint differed as a function of age: juvenile stress produced substantial increases in play fighting, whereas adolescent and adult stress resulted in decreases in social investigation and social preference. The KOR agonist U62066 dose-dependently reduced social behaviors in non-stressed adults, producing social avoidance at the highest dose tested, while younger animals displayed reduced sensitivity to this socially suppressing effect of U62066. Interestingly, in stressed animals, the socially suppressing effects of the KOR agonist were blunted at all ages, with juveniles and adolescents exhibiting increased social preference in response to certain doses of U62066. Taken together, these findings support the hypothesis that the DYN/KOR system changes with age and differentially responds and adapts to stress across development.

## Introduction

The dynorphin/kappa opioid receptor (DYN/KOR) system has been identified as a potential target for the treatment of various disorders associated with stress, including anxiety, depression, and alcohol/substance use disorders (Anderson & Becker, 2017; Chavkin & Ehrich, 2014; Crowley & Kash, 2015; Knoll & Carlezon, 2010; Schwarzer, 2009; Tejeda, Shippenberg, & Henriksson, 2012; Van’t Veer & Carlezon, 2013). Numerous studies in both humans and animals have demonstrated that activation of the DYN/KOR system is associated with behavioral alterations, including aversion, dysphoria, and anxiety, that resemble the effects of stress (Hang, Wang, He, & Liu, 2015; Van’t Veer & Carlezon, 2013). Additionally, blockade of DYN/KOR signaling, either through administration of KOR antagonists or through genetic or viral downregulation/knock-down of DYN and/or KORs, attenuates stress-associated behavioral changes (McLaughlin, Li, Valdez, Chavkin, & Chavkin, 2006; McLaughlin, Marton-Popovici, & Chavkin, 2003) and molecular adaptations within various brain structures known to be affected by stress (Bruchas & Chavkin, 2010; Crowley & Kash, 2015; Lemos et al., 2012).

Despite an extensive literature demonstrating dysphoric/aversive effects following activation of the DYN/KOR system, several studies have reported either opposite effects or a lack of response/sensitivity to KOR agonists (Hang et al., 2015). For example, systemic administration of KOR agonists produced anxiolytic effects on the elevated plus maze (Alexeeva, Nazarova, & Sudakov, 2012; Braida et al., 2009; Kudryavtseva, Gerrits, Avgustinovich, Tenditnik, & Van Ree, 2006; Privette & Terrian, 1995). Similarly, microinjections of a KOR agonist into the infralimbic cortex also produced an anxiolytic effect (Wall & Messier, 2000b), while microinjection of the KOR antagonist, nor-BNI, resulted in anxiogenesis (Wall & Messier, 2000a). While these paradoxical effects of manipulations of the DYN/KOR system have been largely attributed to procedural and methodological differences across studies, a common factor that these studies share is that animals were tested early in life (Diaz, Przybysz, & Rouzer, in press). Surprisingly few studies, however, have directly compared responses to manipulations of the DYN/KOR system between younger and older animals. For instance, adolescent rats were found to be insensitive to conditioned place aversion associated with systemic administration of a KOR agonist, an effect that was observed in adult rats (Anderson, Morales, Spear, & Varlinskaya, 2014). Another study reported that KOR activation in pre-weanlings increased appetitive responding for water (Petrov, Nizhnikov, Varlinskaya, & Spear, 2006), opposite to what had been shown in adults (Bals-Kubik, Herz, & Shippenberg, 1989). Moreover, a selective KOR antagonist, norbinaltorphimine (nor-BNI), attenuated ethanol-induced taste aversion only in stressed adults, with stressed adolescents being insensitive to the effects of nor-BNI (Anderson, Agoglia, Morales, Varlinskaya, & Spear, 2013). In support of these notably different behavioral effects associated with pharmacological activation and/or suppression of the DYN/KOR system across ontogeny, we recently found that KOR activation in the basolateral amygdala (BLA) of adolescent rats increased GABA transmission, without having an effect in the adult BLA (Przybysz, Werner, & Diaz, 2017). An age-dependent increase followed by a decrease in DYN-mediated hyperpolarization of neurons within the paraventricular nucleus of the thalamus has also been shown (Chen et al., 2015). Hence, there is clear behavioral and cellular evidence consistent with age-dependent differences in the functional role of the DYN/KOR system (Diaz et al., in press).

Although the DYN/KOR system has been demonstrated to be both engaged in and altered by stress, resulting in enhanced anxiety (Crowley & Kash, 2015; Schwarzer, 2009; Tejeda et al., 2012; Van’t Veer & Carlezon, 2013), the influence of age and sex has not been carefully examined. Anxiety-like behavior in rats has been extensively assessed using the social interaction test [see (File & Seth, 2003) for references and review]. In the conventional social interaction test, a pair of rats is placed into a testing arena, and overall time spent in social interactions is generally used as a dependent measure (File & Hyde, 1978). However, this approach combines together the discrete behavioral acts (e.g., sniffing of a partner, social grooming, following, chasing, pinning, etc.) that reflect behaviorally distinctive and differentially regulated forms of social behavior, including social investigation and play fighting. These social behaviors are characterized by distinguishable developmental patterns (Vanderschuren, Niesink, & Van Ree, 1997; Varlinskaya & Spear, 2008; Varlinskaya, Spear, & Spear, 1999) and differential responsiveness to anxiety-provoking manipulations (Doremus-Fitzwater, Varlinskaya, & Spear, 2009b). For example, play fighting shows an inverted U-shaped ontogenetic pattern, peaking around postnatal day (P) 30-35. In contrast, social investigation increases with age, representing a more adult-typical form of social behavior (Vanderschuren et al., 1997; Varlinskaya et al., 1999). Play fighting, but not social investigation, is drastically increased by isolate housing throughout the juvenile and adolescent periods (Vanderschuren et al., 1997; Varlinskaya & Spear, 2008), while social investigation is exclusively decreased by prior history of exposure to non-social stressors during adolescence and in adulthood (Doremus-Fitzwater et al., 2009b; Varlinskaya, Doremus-Fitzwater, & Spear, 2010). Together, these findings suggest that play fighting and social investigation may be differentially affected by pharmacological activation of the DYN/KOR system. Our modification of the social interaction test (Varlinskaya et al., 1999) allows an experimental animal to freely move toward or away from a non-manipulated social partner in a 2-compartment testing apparatus, thereby permitting assessment of social preference and/or avoidance in addition to measurement of the frequencies of play fighting and social investigation (Varlinskaya et al., 1999). Using this modified social interaction test, we have found decreases in social preference and/or social investigation to reflect anxiety-like alterations (Doremus-Fitzwater et al., 2009b; Morales, Varlinskaya, & Spear, 2013; Varlinskaya et al., 2010; Varlinskaya & Spear, 2012).

Given the mounting evidence demonstrating age-related differences in vulnerability to and outcomes of stress exposure (Enoch, 2011; Romeo, 2017; Tottenham & Galvan, 2016) as well as in responsiveness to the aversive effects of the DYN/KOR system activation (Anderson et al., 2014), the present study was designed to systemically assess the effects of pharmacological activation of the DYN/KOR system on social investigation, social preference, and play fighting across ontogeny in non-stressed males and females as well as males and females with a prior history of repeated exposure to restraint.

## Methods

### Subjects

Juvenile, adolescent and adult Sprague-Dawley male and female rats bred and reared in our colony at Binghamton University were used. A total of 72 litters provided 360 male and female offspring to serve as experimental subjects and 360 to serve as partners. Animals were housed in a temperature-controlled (22°C) vivarium, and maintained on a 12:12 hr light:dark cycle (lights on at 0700 hr) with ad libitum access to food (Purina rat chow) and water. Litters were culled to 10 pups (five males and five females) within 24 hr after birth on P0 and reared until weaning with their mothers in standard plastic maternity cages with pine shavings as bedding material. Rats were weaned on P21 and housed with their same-sex littermates. At all times, rats used in the current study were produced, maintained, and treated in accordance with the guidelines for animal care established by the National Institutes of Health, using protocols approved by the Binghamton University Institutional Animal Care and Use Committee.

### Experimental Design

The design was a 3 (age: juvenile, adolescent, adult) X 2 (sex) X 2 (stress condition: no stress or repeated restraint) x 5 (U62,066 dose: 0, 0.1, 0.2, 0,3, and 0.4 mg/kg) factorial, with six experimental animals tested per group. Juveniles were tested on P28, adolescents -on P35, and adults were tested on P70. All animals from a given litter were assigned to the same stress condition. To avoid the possible confounding of litter with the experimental variables (Holson & Pearce, 1992; Zorrilla, 1997), no more than one subject per sex from a litter was assigned to a particular U62066 dose/stress condition, with order of testing counterbalanced across litters.

### Stressor Procedures

Beginning at P24 for juveniles, at P30 for adolescents, and at P66 for adults, rats from the repeated stress group were removed from their home cage between 1000 – 1200 hr and then restrained in an age size-adjusted (5.08 cm diameter x 12.7 cm length for juveniles, 6.35 cm diameter x 15.24 cm length for adolescents, and 8.26 cm diameter x 20.32 cm length for adults) flat-bottom restrainers (Braintree Scientific, Braintree, MA) for 90 min in a novel holding cage. For animals in the stress group, this restraint procedure was repeated each day for 5 days. Animals placed in the control condition were non-manipulated throughout the 5-day stressor phase until the time of testing.

As in our previous studies (Doremus-Fitzwater, Varlinskaya, & Spear, 2009a; Varlinskaya et al., 2010; Varlinskaya & Spear, 2012), restraint was used as the stressor, since this stressor is primarily psychological in nature and does not induce physical pain or harm to the experimental subjects (Herman & Cullinan, 1997; Weinberg, Girotti, & Spencer, 2007).

### Drug administration

The selective kappa agonist U62066 (Sigma-Aldrich) was dissolved in 0.9% saline vehicle and injected subcutaneously in a volume of 2 ml/kg 30 min prior to testing. Five doses of the drug were tested: 0 (saline), 0.1, 0.2, 0.3, and 0.4 mg/kg.

### Testing Procedures

Immediately after the 90-min stressor exposure on day 5 (or upon removal from the home cage for non-stressed animals), each subject was injected with one of the five doses of U62066. Immediately after drug administration, each experimental animal was placed alone into a testing chamber for 30 min. This pretest familiarization was conducted to increase baseline levels of social interaction during testing, hence making potential anxiogenic effects of the repeated stressors easier to observe (File & Seth, 2003). A same age and sex test partner unfamiliar with both the test apparatus and the experimental animal was then placed into the apparatus, and social interactions were recorded for 10 min. Partners were always non-stressed, drug-naive animals that had not been socially isolated prior to testing. Weight differences between test subjects and their partners were minimized as much as possible, with this weight difference not exceeding 5 g for animals at P28, 10 g at P35, and 20 g at P70, with test subjects always being heavier than their partners.

Testing was conducted in Plexiglas test chambers (30 x 20 x 20 cm for juveniles and adolescents; 45 x 30 x 30 cm for adults) that contained clean pine shavings. The test apparatuses (Binghamton Plate Glass, Binghamton, NY) were divided into two compartments by a clear Plexiglas partition containing an aperture (7 x 5 cm for juveniles and adolescents; 9 x 7 for adults) to allow movement of animals between compartments (Varlinskaya et al., 1999; Varlinskaya, Spear, & Spear, 2001). Each 10-min social interaction test session was conducted under dim light (15-20 lux) between 1000 and 1400 hr, with a white noise generator used to attenuate extraneous sounds during testing. The behavior of each pair was recorded by a video camera mounted above the apparatus.

### Behavioral Measures

The frequencies of social investigation and play fighting were analyzed from video recordings (Meaney & Stewart, 1981; Thor & Holloway, 1984; Varlinskaya & Spear, 2008) by a trained experimenter without knowledge of the experimental condition of any given animal. Social investigation was defined as the sniffing of any part of the body of the partner. Play fighting was scored as the sum of the frequencies of the following behaviors: pouncing or playful nape attack (experimental subject lunges at the partner with its forepaws extended outward); following and chasing (experimental animal rapidly pursues the partner); and pinning (the experimental subject stands over the exposed ventral area of the partner, pressing the animal against the floor). Play fighting can be distinguished from serious fighting in the laboratory rat by the target of the attack—during play fighting, snout or oral contact is directed towards the partner’s nape, whereas during serious fighting the partner’s rump is targeted (Pellis & Pellis, 1987). Aggressive behavior (serious fighting) was not analyzed in these experiments, since subjects did not exhibit serious attacks or threats.

Social preference/avoidance was assessed by separately measuring the number of crossovers from one side of the apparatus to the other demonstrated by the experimental subject towards as well as away from the social partner and was indexed by means of a coefficient of preference/avoidance [coefficient (%) = (crossovers to the partner – crossovers away from the partner)/(total number of crosses both to and away from the partner) x 100]. Social preference was defined as positive values of the coefficient, while social avoidance was associated with negative values (Varlinskaya et al., 1999).

The total number of crossovers (movements between compartments through the aperture to and from the social partner) exhibited by each experimental subject was used as an index of locomotor activity in the social context (Varlinskaya et al., 1999).

### Data Analyses

Data for each dependent variable (play fighting, social investigation, preference coefficient, and total number of crossovers) were analyzed using separate 3 (age) X 2 (sex) X 2 (stress condition) X 5 (U62,066 dose) ANOVAs. In order to avoid inflating the possibility of type II errors on tests with at least three factors (Carmer, 1973), Fisher’s planned pairwise comparison test was used to explore significant effects and interactions. Where significant interactions involving stress condition and U62066 challenge dose were evident, drug-induced changes were assessed between U62066-challenged animals and saline-challenged controls within each stress condition separately for each age as well as between non-stressed and stressed animals of the same age at a certain dose of U62066. Sensitivity to the effects of U62066 was assessed by the lowest dose that produced significant changes relative to saline within each age/stress condition.

## Results

### Social Investigation

The significant Sex X Drug Dose interaction, F (4, 300) = 2.78, p < 0.05 reflected a general decrease in sensitivity of females to U62066-induced suppression of social investigation relative to males. Females exhibited significant decreases at the doses of 0.2, 0.3, and 0.4 mg/kg, whereas in males all four doses produced a significant reduction of social investigation (see Table 1). Significant sex differences were also evident at the doses of 0.3 and 0.4 mg/kg, with females being less affected than males.

The ANOVA of social investigation also revealed a significant Age X Stress X U62066 dose interaction, F(8, 300) = 3.39, p < 0.001. When data were collapsed across sex to examine these interactions, restraint was found to significantly suppress baseline social investigation (i.e., levels of social investigation seen after saline injection) in adolescent and adult animals only, with no socially suppressing effects of juvenile stress evident in young animals **(Fig. 1)**. Assessment of age-related differences in sensitivity to the socially suppressing effects of the kappa opioid agonist in non-stressed animals revealed significant dose-dependent decreases in social investigation at all doses of U62066 in juveniles and adults, whereas in adolescents the minimal effective dose was 0.3 mg/kg **(Fig. 1)**. Stress-induced changes in sensitivity to the kappa agonist also differed as a function of age. Stressed animals became less sensitive to U62066-induced suppression of social investigation, with this stress effect being more pronounced in adolescents and adults than in juveniles. Specifically, in stressed juveniles, social investigation was significantly reduced at all doses except for the lowest dose of 0.1 mg/kg. In contrast, stressed adolescents became substantially less sensitive to the social suppression induced by U62066 and responded by a significant decrease in social investigation only to the highest dose of 0.4 mg/kg. Similarly, stressed adult animals demonstrated significant suppression of social investigation at and 0.4 mg/kg of the kappa agonist, with no significant decreases emerging at lower doses.

**Figure 1.**
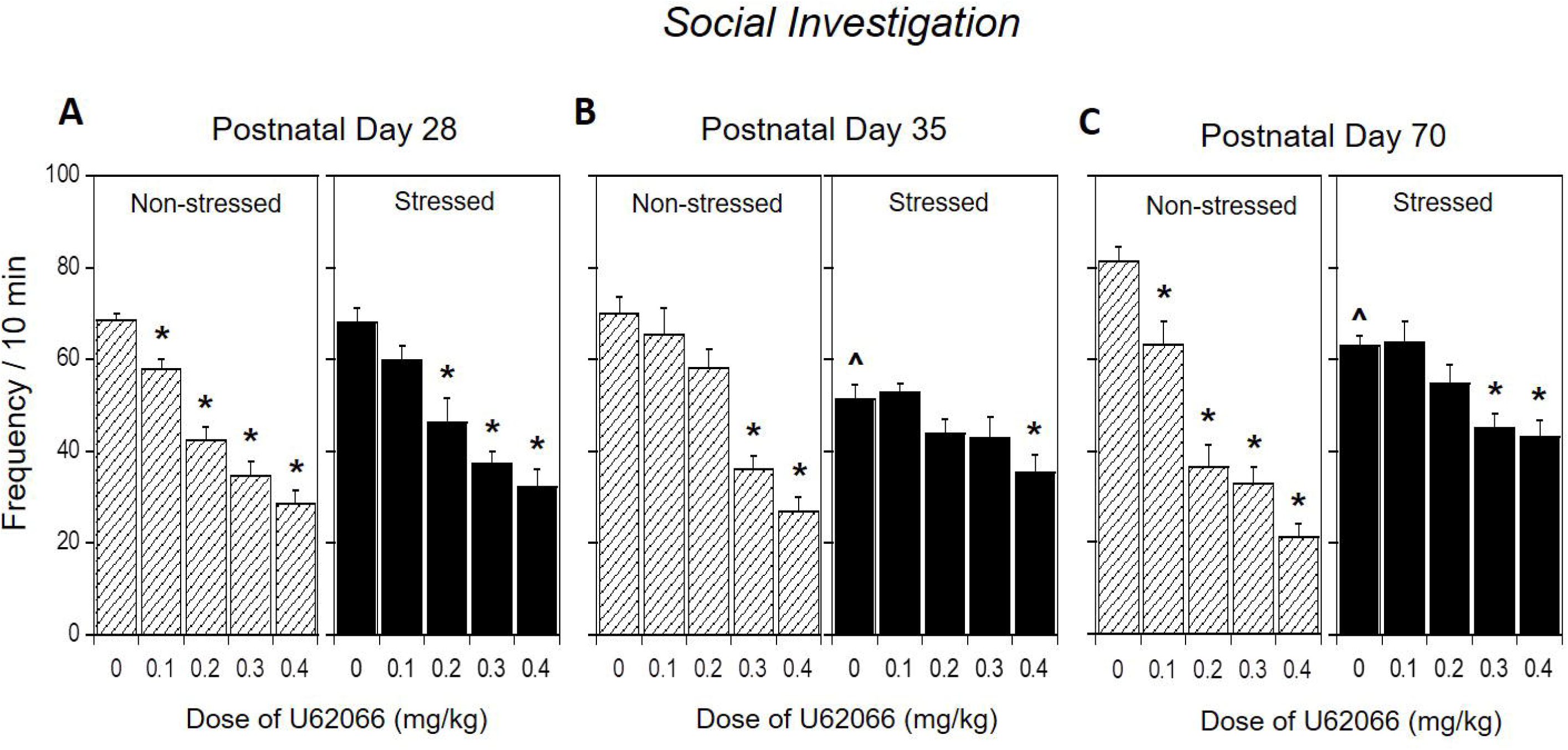
Social preference/avoidance in stressed and non-stressed rats: Effects of the selective KOR agonist U62066. Asterisks (*) denote significant drug effects within each age/stress exposure condition relative to vehicle. (^) denote significant stress-related changes in each age group under basal (0 mg/kg dose) conditions. Data are expressed as mean ± SEM, p < 0.05 and collapsed across sex.

### Social Preference/Avoidance

The ANOVA revealed a significant Age x Stress X U62066 dose interaction, F(8, 300) = 2.33, p < 0.05. Baseline social preference again differed as a function of stress conditions in adolescent and adult, but not juvenile, rats, with exposure to the stressor significantly decreasing social preference relative to non-stressed animals among adolescents and adults but not among stressed juveniles. Marked age differences in the coefficient were evident in non-stressed animals (**Fig. 2**). Non-stressed juveniles were less sensitive to the increase in social anxiety-like effects of U62066 than older animals, showing significant decreases in social preference at the highest dose only, whereas non-stressed adolescents demonstrated significant decreases in social preferences at the doses of 0.3 and 0.4 mg/kg. Non-stressed adults were the most sensitive to the anxiogenic effects of U62066, showing negative values of the coefficient at 0.2, 0.3, and mg/kg doses, with the highest dose of this kappa agonist (0.4 mg/kg) producing substantial social avoidance in adults. These age-dependent anxiogenic effects of the selective kappa agonist were eliminated by repeated restraint, with no decreases in social preference relative to corresponding saline-injected controls evident at either age. Interestingly and in contrast, juveniles showed significant increases in social preference at 0.4 mg/kg, with a similar socially anxiolytic effect evident in adolescents at the lowest dose of 0.1 mg/kg.

**Figure 2.**
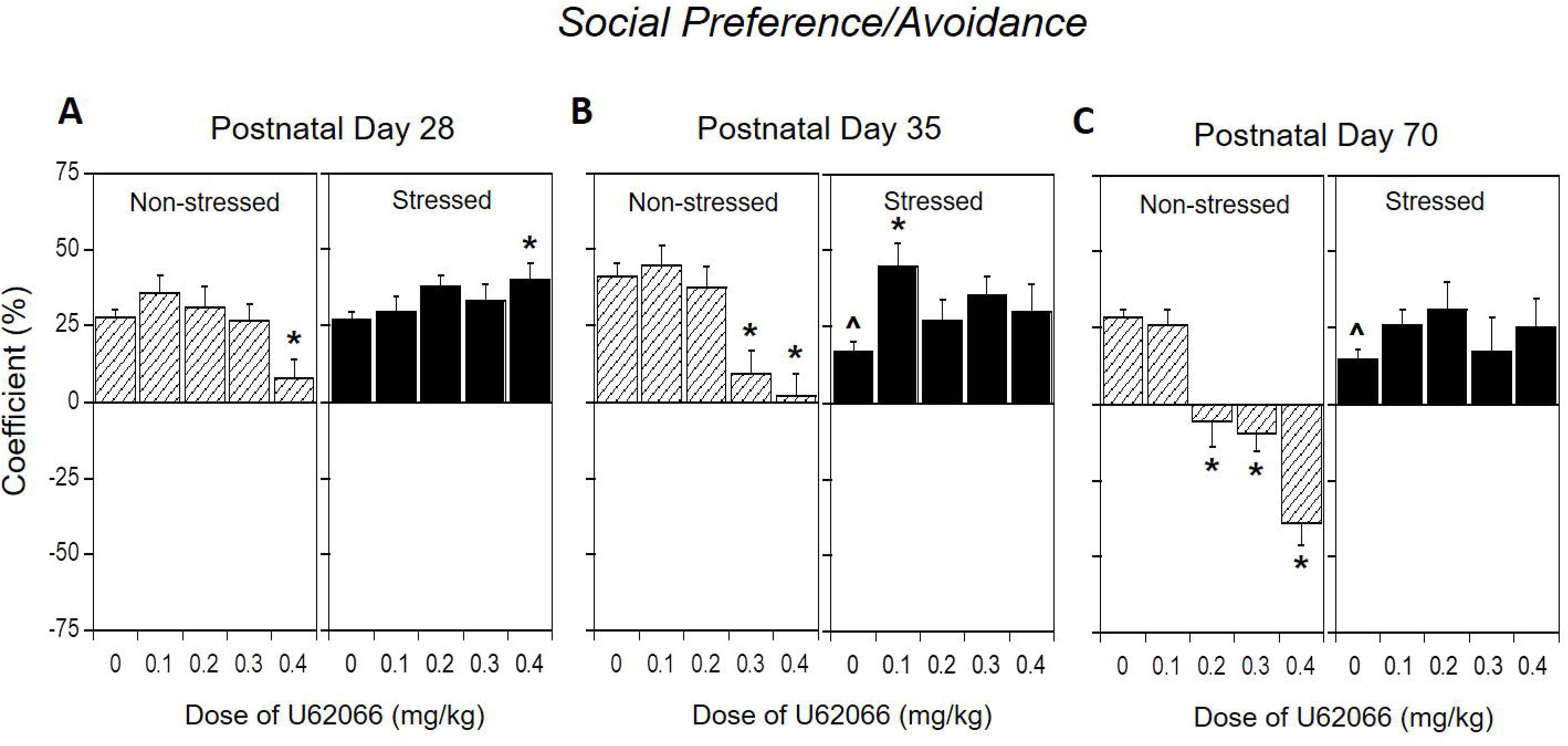
Social investigation in stressed and non-stressed rats: Effects of the selective KOR agonist U62066. Asterisks (*) denote significant drug effects within each age/stress exposure condition relative to vehicle. (^) denote significant stress-related changes in each age group under basal (0 mg/kg dose) conditions. Data are expressed as mean ± SEM, p < 0.05 and collapsed across sex.

### Play Behavior

A significant Age x Stress X U62066 dose interaction, F(8, 300) = 2.84, p < 0.01, was evident for play behavior. The most striking effect of stressor exposure was evident in juveniles, with exposure to restraint markedly increasing play behavior under basal, saline challenge conditions. In non-stressed animals all four doses of U62066 significantly suppressed play behavior regardless of age (**Fig. 3**). In stressed adolescents and adults the suppressing effects of the lowest dose did not reach statistical significance.

**Figure 3.**
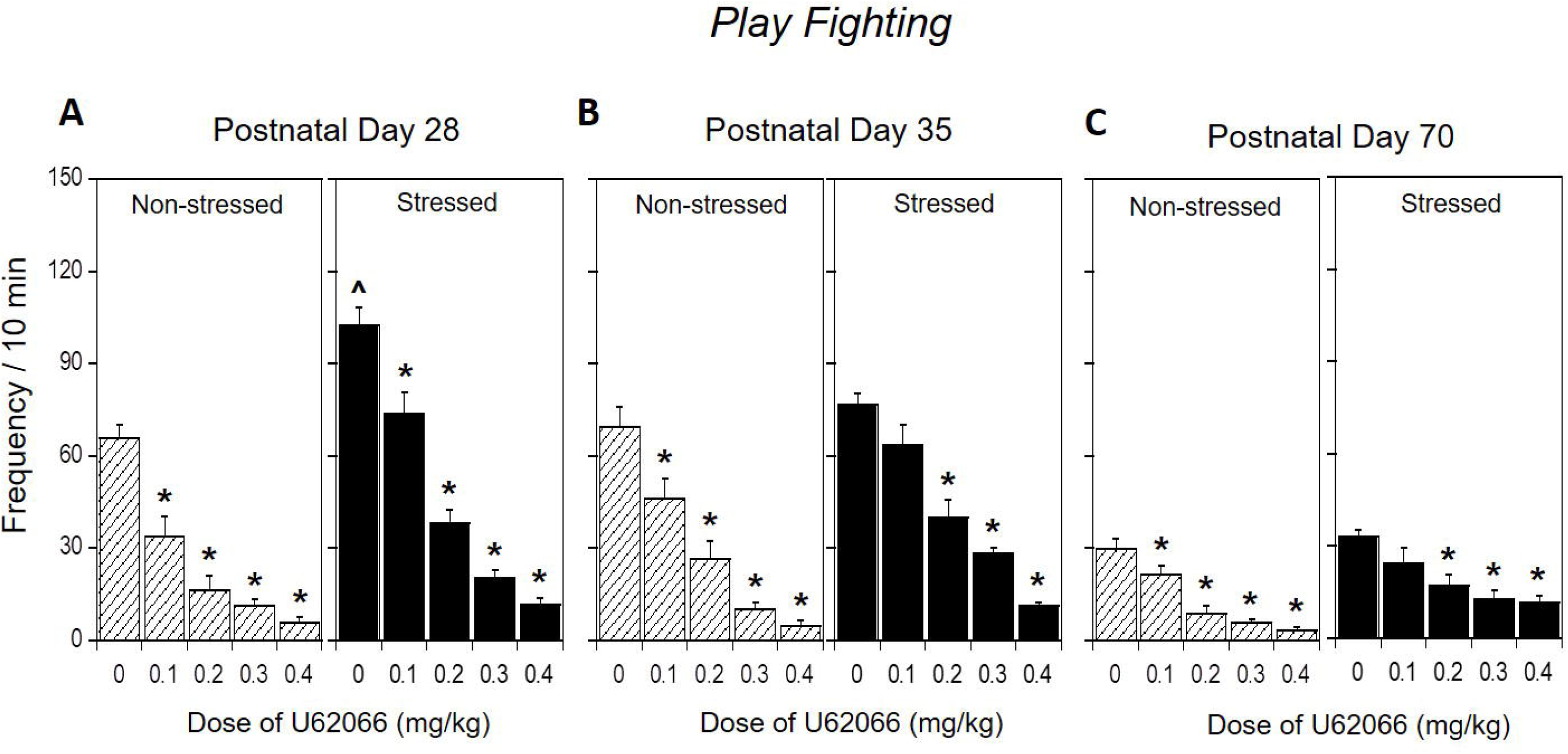
Play fighting in stressed and non-stressed rats: Effects of the selective KOR agonist U62066. Asterisks (*) denote significant drug effects within each age/stress exposure condition relative to vehicle. (^) denote significant stress-related changes in each age group under basal (0 mg/kg dose) conditions. Data are expressed as mean ± SEM, p < 0.05 and collapsed across sex.

### Locomotor Activity (Table 2)

The analysis of general locomotor activity under social circumstances indexed via total number of crossovers between compartments revealed significant main effects of age, F (2, 300) = 25.39, p < 0.0001 and drug dose, F (4, 300) = 143.27, p < 0.0001. In general, juveniles and adolescents demonstrated significantly more crossovers than their older counterparts, with all doses of U62066 gradually suppressing locomotor activity under social test circumstances regardless of age or stress condition (**Table 2**).

## Discussion

Given evidence that the DYN/KOR system in younger animals may function and adapt to stressors differently early in life than adulthood [see review by (Diaz et al., in press)], in this study we sought to systematically test the effects of a selective KOR agonist on social investigation, social preference, and play fighting in juveniles, adolescents and adults. We also examined the impact of repeated restraint on responsiveness to KOR activation and assessed whether sex differences existed in these measures. In non-stressed animals, pharmacological activation of KORs suppressed social behavior at all ages, although age differences in sensitivity to this social suppression varied with the social measure under investigation. Repeated restraint suppressed social behaviors in adolescent and adult animals, while conversely notably *increasing* social play among juveniles. Stress also generally decreased the response to U62066 at all ages, effects that were particularly notable for social preference/avoidance in mature animals, with U-62066-induced social avoidance in non-stressed adults converted to social preference following repeated restraint stress. Finally, we also found that in general, females were less sensitive to the effects of KOR activation on social investigation, but not other social measures, when collapsed across age and stress condition.

### Ontogenetic social effects of U62066 in non-stressed animals

It is well established that activation of the DYN/KOR system results in dysphoria, aversion, and increased anxiety-like behaviors in both humans and animals (Hang et al., 2015; Schwarzer, 2009; Van’t Veer & Carlezon, 2013). However, numerous studies have reported opposite effects or insensitivity to KOR activation during early ontogeny (Diaz et al., in press). For instance, neonates demonstrated increased appetitive responding to a surrogate nipple providing water following administration of a KOR agonist (Petrov et al., 2006). A different study found that adolescents were insensitive to KOR activation assessed in a conditioned place preference/aversion paradigm, whereas adults showed conditioned place aversion to the same doses of a selective KOR agonist (Anderson et al., 2014). Several studies have also reported anxiolytic effects of KOR agonists on the elevated plus-maze in rats that were tested at a weight consistent with adolescence (∼150-210 grams) (Alexeeva et al., 2012; Braida et al., 2009; Privette & Terrian, 1995). Taken together, there is strong behavioral evidence that the DYN/KOR system functions differently across ontogeny. Consistent with these behavioral observations, we found that adolescents were less sensitive to the reductions in social investigation induced by U62066 relative to both juveniles and adults. However, a gradual ontogenetic increase in responsiveness to U62066 was evident when social preference was the measure under investigation, with P28 animals being the least sensitive and adults being the most sensitive to U62066-induced decreases in social preference. Adults not only demonstrated significant reductions in social preference at lower doses, but also exhibited social avoidance at the highest dose (0.4 mg/kg), indicating that this dose was severely anxiogenic to that age group. Overall, these findings complement and add to the growing literature demonstrating that animals are less sensitive to the dysphoric, aversive, and anxiogenic effects of KOR agonists during ontogeny.

While the mechanism(s) underlying this early-life insensitivity are unknown, we previously found that activation of KORs in the BLA increases GABA transmission in adolescent males, but not in adult males, while not affecting glutamate transmission at either age (Przybysz et al., 2017). This would result in reduced excitability of the BLA, which has been shown following KOR activation in younger animals (Huge, Rammes, Beyer, Zieglgansberger, & Azad, 2009), and ultimately reduced social (Truitt et al., 2007) and non-social anxiety-like behaviors (Bueno, Zangrossi, & Viana, 2005; Lack, Diaz, Chappell, DuBois, & McCool, 2007). In contrast, we also found that KOR activation reduces GABA transmission in the central amygdala of both adolescent and adult males (Przybysz et al., 2017). Interestingly, another study found an age-dependent change in KOR-induced outward currents in the paraventricular nucleus of the thalamus, a brain structure that interfaces with the anxiety circuit, whereby the magnitude of these currents increases between 2 and 4 weeks of age, followed by a reduction by 8 weeks of age (Chen et al., 2015). Based on these various ontogenetic differences in KOR function in brain structures associated with anxiety, it is possible that systemic administration of a KOR agonist engages potentially opposing actions, resulting in no net effect, purportedly observed as an insensitivity to the drug. However, the current study suggests that the dose-response curve for the selective KOR agonist is shifted to the right in younger animals relative to adults, as the highest doses produced increases in social anxiety-like behavior even in younger animals. Studies examining these mechanisms are warranted and are currently underway.

### Stress differentially affects social behavior across ontogeny

Our general knowledge of age-dependent differences in the expression of various behaviors is extensive, and there is compelling evidence suggesting that the impact of stress in early-life may be more detrimental due, in part, to alterations in critical neurodevelopmental processes (Romeo, 2017; Tottenham & Galvan, 2016). Consistent with this, in our previous studies, adolescents and adults showed similar patterns of social alterations associated with repeated restraint (Varlinskaya et al., 2010), whereas juveniles differed drastically from older animals (Varlinskaya, Truxell, & Spear, 2013). Similarly, in the present study, adolescents and adults demonstrated significant stress-related decreases in social investigation and social preference that were not evident in juveniles. Instead, juveniles responded to the prior stress exposure with an enhancement of play fighting – a form of social behavior that is more expressed in juveniles and adolescents than in adults (Vanderschuren et al., 1997; Varlinskaya et al., 1999). These observed stress-associated alterations were rather specific for social behavior, since overall locomotor activity, indexed via total number of crossovers, was not affected by stress at either age. One possible explanation for these age-related differences in social responsiveness to repeated restraint is that juveniles tested at P28 do not respond to stressful and hence anxiety-provoking manipulations in a way their older counterparts do. This explanation is unlikely, however, since juvenile rats, similar to their more mature counterparts, respond to a novel, anxiety-provoking test situation by enhanced social anxiety-like behavior, indexed by transformation of social preference into social avoidance (Varlinskaya & Spear, 2006, 2008). An alternative possibility is that the 90-min periods of restraint were perceived by juveniles as significant social deprivation, with this social deprivation producing substantial increases in play fighting (Varlinskaya & Spear, 2008). Timing of assessment following stress is clearly an important aspect to consider, since different adaptations occur in the DYN/KOR over time (Knoll & Carlezon, 2010). Studies examining these time-course-dependent adaptations across ontogeny are ongoing in our lab and will shed light on these factors.

Some researchers suggest that pre-pubertal animals differ markedly in their responsiveness to stress from post-pubertal, adult rats (Koenig, Walker, Romeo, & Lupien, 2011; McCormick, Merrick, Secen, & Helmreich, 2007; Romeo, 2010). Pre-pubertal stress has been shown to have long-lasting consequences. For instance, alterations in stress responsiveness in adulthood were evident following even a brief, acute stress exposure on P28 (Avital & Richter-Levin, 2005). Furthermore, pre-pubertal stress enhanced anxiety-like behavior and substantially reduced exploratory behavior in adulthood (Jacobson-Pick & Richter-Levin, 2010; Tsoory, Cohen, & Richter-Levin, 2007). However, when these young animals were tested immediately after exposure to stressors, they demonstrated increases in exploratory behavior (Horovitz, Tsoory, Hall, Jacobson-Pick, & Richter-Levin, 2012) and decreases in anxiety (Jacobson-Pick & Richter-Levin, 2012), findings that are reminiscent of the results of the present study. Therefore, it is possible that the observed age differences in the social consequences of exposure to restraint are related, to some extent, to pubertal maturation. This possibility seems unlikely, however, in that adolescent male rats tested at P35, while still pre-pubertal (Vetter-O’Hagen & Spear, 2012), demonstrated adult-like social responding to repeated restraint. The main difference between the two younger groups in the present study was that they were exposed to the stressor either as juveniles or as adolescents. Together, these reports along with the findings from the current study suggest that the drastic differences in the social consequences of repeated restraint between juveniles and adolescents may not be puberty-related but rather associated with age-related neural alterations (Spear, 2000, 2011). Clearly more studies are required for better understanding immediate and long-lasting consequences of juvenile versus adolescent stress.

### Stress differentially alters sensitivity to KOR activation across ontogeny

One of the primary objectives of this study was to determine how stress alters the effects of pharmacological activation of KORs across ontogeny. In general, stress diminished responses to U62066 with these effects being least pronounced among juveniles, and most marked in adults. Specifically, doses of the KOR agonist that produced robust reductions in social investigation and social preference in non-stressed adolescents and adults no longer produced those effects following restraint. In fact, in adolescents, the lowest dose of U62066 (0.1mg/kg) reversed the socially anxiogenic effect of stress indexed via significant reductions in social preference, demonstrating a potentially anxiolytic effect of KOR activation at this low dose. A similar effect of the highest dose of U62066 (0.4mg/kg) was evident in stressed juveniles, with this dose significantly increasing social preference relative to vehicle. While the mechanisms driving these alterations in the DYN/KOR system are not clear, the diminished effects of U62066 on social investigation and social preference (see Figures 1 and 2) are likely a result of the DYN/KOR system already being activated and potentially saturated during the test, given that testing was performed almost immediately following the final restraint. In contrast, while the anxiolytic effects are difficult to interpret, studies have reported anxiolytic effects of KOR agonists in younger animals. Interestingly, studies reporting such anxiolytic effects have used animals that were likely or confirmed to have been ordered and shipped from an animal vendor (Alexeeva et al., 2012; Braida et al., 2009; Privette & Terrian, 1995). There is compelling evidence that shipping causes major stress in animals, which is mimicked by adolescent social isolation, and produces numerous behavioral and physiological alterations (Chappell, Carter, McCool, & Weiner, 2013; Rau, Chappell, Butler, Ariwodola, & Weiner, 2015). Therefore, these data suggest that early-life stress can result in adaptations in the DYN/KOR system that produce opposite effects to those seen in the adult DYN/KOR system. Studies investigating the underlying mechanisms are needed to better understand these age-dependent neuroadaptations to the DYN/KOR system.

### Age, stress and DYN/KOR system’s roles in play behavior

Adolescent and adult animals demonstrated substantial reductions in sensitivity to effects of U62066 on social preference, and to some extent, social investigation --two social behaviors previously shown to be stressor-sensitive (Varlinskaya et al., 2010; Varlinskaya et al., 2013). Yet, more limited stress-associated changes were evident in all age groups when play fighting was the measure under investigation. In general, play fighting demonstrated by juvenile and adolescent rats is considered to be under control of the endogenous opioid systems (Trezza, Baarendse, & Vanderschuren, 2010). Pharmacological activation of mu opioid receptors enhances play fighting in juvenile and adolescent rats [reviewed in (Trezza et al., 2010)], whereas pharmacological activation of KORs decreases social play behavior (Vanderschuren, Niesink, Spruijt, & Van Ree, 1995). The results of the present study are in agreement with these early findings in that the selective KOR agonist effectively and dose-dependently decreased play fighting in adolescent and adult animals. However, it seems unlikely that the endogenous DYN/KOR system is involved in modulation of play fighting in previously stressed animals, given that no baseline decreases in play fighting were seen in saline control animals in response to stress-induced activation of this system. In contrast, stress exposure substantially increased play fighting in juveniles. Taken together with the age-specific effects of repeated restraint on social investigation and social preference, it is likely that the endogenous DYN/KOR system plays little role if any in modulation of play behavior in previously stressed animals.

### Conclusions

The present study showed that in non-stressed animals, age differences in sensitivity to the socially suppressing effects of the selective KOR agonist, U62066, strongly depend on the measure under investigation, with the most pronounced ontogenetic increases evident in sensitivity to agonist-induced social avoidance. Findings of this study also demonstrated that responsiveness to repeated restraint stress in terms of both stress-induced behavioral alterations and stress-associated changes in sensitivity to the social consequences of pharmacological activation of the DYN/KOR system differs notably in juveniles relative to adolescents and adults. Consistent with our previous findings, stress-induced suppression of social investigation and social preference were evident in adolescents and adults, but not juveniles, with this youngest age group instead demonstrating substantial stress-induced increases in play fighting (Doremus-Fitzwater et al., 2009b; Doremus-Fitzwater, Varlinskaya, & Spear, 2010). Furthermore, more robust stress-associated decreases in responsiveness to pharmacological KOR activation were seen in adults, and to some extent adolescents, than juveniles, suggesting ontogenetic increases in stress-induced activation of the DYN/KOR system.

In summary, this study provides compelling evidence supporting the under-studied phenomenon of ontogenetic changes in the native DYN/KOR system, particularly demonstrating that stress-related alterations in the DYN/KOR system function are age-dependent. Given that the DYN/KOR system has been suggested to be a potential pharmacological target for stress-related disorders, such as anxiety and addiction (Chavkin & Koob, 2016; Tejeda et al., 2012), this study suggests that the age and timing of stress exposure must be taken into consideration.

## Acknowledgements

Research reported in this publication was supported by the National Institute on Alcohol Abuse and Alcoholism of the National Institutes of Health under Award Numbers P50AA017823 and R03AA024890.

